# Distance-dependent changes in courtship song amplitude reflect song state changes

**DOI:** 10.1101/277210

**Authors:** Harini Suri, Raghav Rajan

## Abstract

Many animals increase the amplitude of their vocalizations as distance from a receiver increases. However, behavioral mechanisms underlying this increase remain unclear. Here, we addressed this using head-attached microphones to accurately record song amplitude in male zebra finches while presenting a female at difference distances. We show that individual courtship song syllables either increased (5/13) or decreased their amplitude (4/13) as distance from the female increased. Both increases and decreases were part of more general acoustic changes that resulted in properties more similar to those of “undirected” songs typically produced in the absence of a female. Increasing distance also reduced female responses to songs and absence of female responses reduced courtship song amplitude and number of songs per bout. These data suggest a simple behavioral mechanism where distance-dependent song amplitude changes reflect song state changes. Such state changes may be a general mechanism underlying distance-dependent amplitude changes in other organisms.

## INTRODUCTION

Animal vocalizations are one form of acoustic communication that serves a multitude of functions including courting mates, sending warning signals about predators, and hunting prey (Bradbury and Vehrencamp, 2011). Such communication is dependent on the distance between sender and receiver as sound degrades with increasing distance (Bradbury and Vehrencamp, 2011; Brenowitz, 1986). Previous studies have shown that many animals produce louder vocalizations as distance from a receiver increases, possibly to compensate for distance-dependent sound degradation (Brenowitz, 1986; Brumm and Slater, 2006; Coen et al., 2016). However, the behavioral mechanisms underlying this increase remain poorly understood.

One well-studied example of a courtship vocalization is adult male zebra finch song (Fee and Scharff, 2010). Song is a stereotyped sequence of vocalizations separated by silent gaps (Fee and Scharff, 2010). Song coupled with a visual display directed towards female birds is used by males for courtship (Sossinka and Böhner, 1980; Ullrich et al., 2016; Williams, 2004). Male zebra finches increase the amplitude of their courtship songs as distance from the female increases (Brumm and Slater, 2006). Brumm and Slater hypothesized that such changes are a result of birds learning that only louder songs elicit responses in females that are far away. However, currently there is no empirical support for this hypothesis.

In addition to courtship song, zebra finches also sing “undirected” song in the absence of the female (Sossinka and Böhner, 1980). Undirected song differs from courtship song in a number of acoustic features (Kao et al., 2005; Sossinka and Böhner, 1980) and is thought to serve as advertisement of courtship potential (Dunn and Zann, 1996a; Dunn and Zann, 1996b). Such advertisements song in other birds are often louder than the courtship songs (Dabelsteen et al., 1998) as they are targeted at birds that are farther away. This suggests an alternate hypothesis to explain distance-dependent song amplitude increases, namely, a change from courtship song to advertisement song properties. This hypothesis makes two specific predictions, namely, (1) undirected song should be louder than courtship song at the shortest distance from the female and (2) other acoustic features of courtship song should become “undirected-like” with increasing distance from the female.

Here, we tested these two predictions by recording and analyzing songs produced by male zebra finches presented with females at different distances. We used head-attached microphones to accurately measure song amplitude independent of the bird’s position within the cage and show that song syllable amplitude and other features of courtship songs became “undirected-like” with increasing distance. Female responses also decreased with distance and absence of female responses reduced song amplitude and the number of songs/bout. These results suggested that distance-dependent song amplitude changes are the result of a more general change to “undirected-like” song properties.

## RESULTS

Songs (Fig. 1A) and visual displays of adult male zebra finches were recorded in the presence of a female placed at one of five distances (Fig. 1B, n=12 birds). They were also recorded in the presence of an empty cage placed at one of the distances (Fig.1C) to characterize undirected song properties. Visual displays associated with all song bouts were screened for previously described characteristics of female-directed courtship songs (Morris, 1954; Ullrich et al., 2016; Williams, 2001) and song bouts categorised as courtship songs were considered for further analysis (see Methods). Consistent with earlier data (Brumm and Slater, 2006), we found a significant decrease in the proportion of courtship song bouts produced by each male as distance from the female increased (Fig. 1D, r = −0.8552, p = 3.07′ 10^−22^).

**FIGURE 1.**
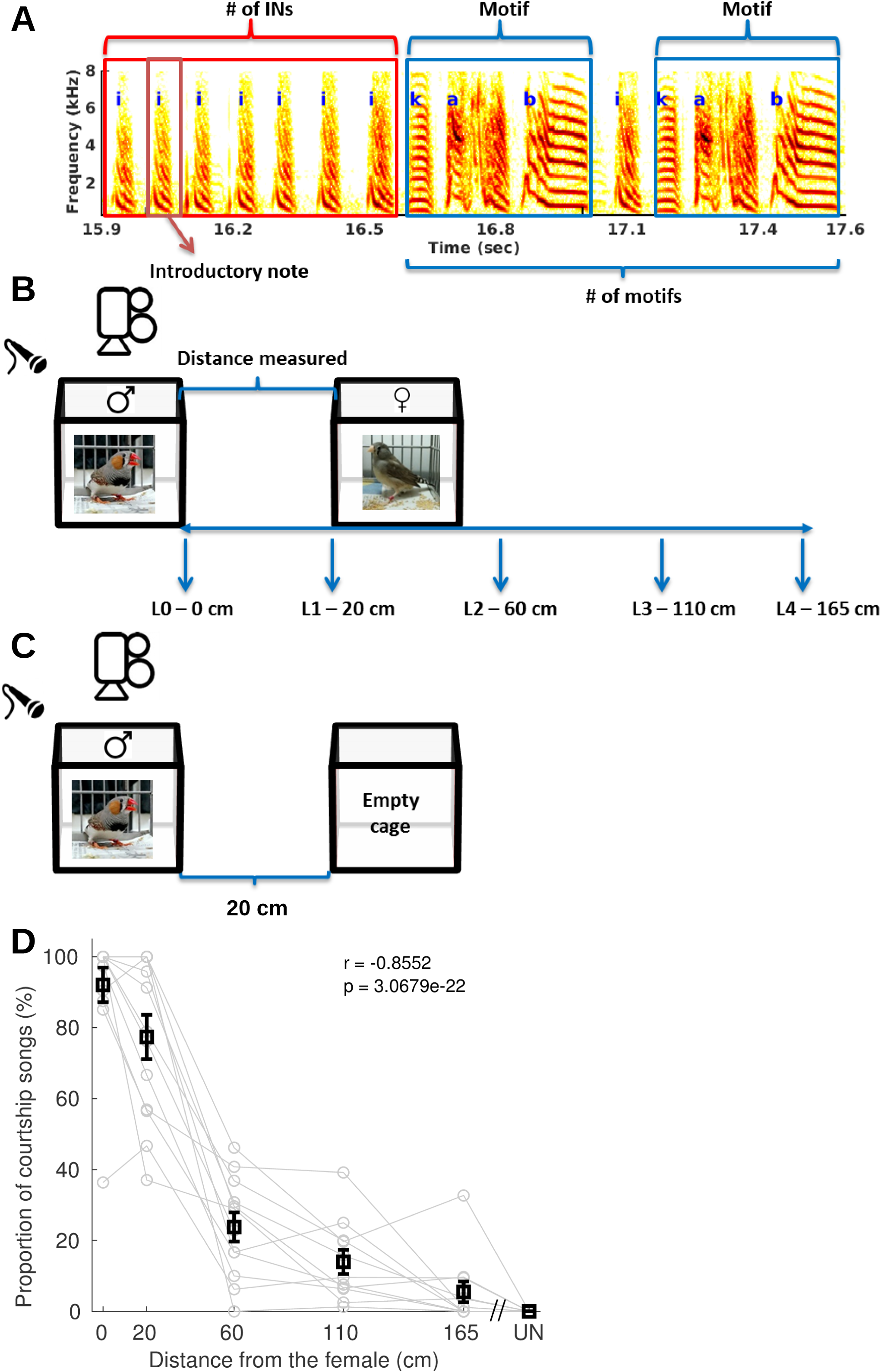
Experimental design and proportion of courtship songs decreased with distance (A) Spectrogram of a song bout beginning with introductory notes (INs) followed by two motifs. ‘i’ represents INs, ‘k’, ‘a’ and ‘b’ represent motif syllables. Three of the song features measured in this study have been shown on this spectrogram - # of INs (red box), # of motifs (blue bracket), and motif durations (blue boxes). (B) Schematic of the experimental setup representing the various distances at which the female bird was presented to the male bird. The male bird’s cage was at a fixed position where we recorded song and behaviour using a microphone and a webcam respectively. (C) The male bird’s song and behaviour were also recorded in the absence of the female, with an empty cage at 20 cm distance from the male bird’s cage. (D) shows changes in the percentage of songs scored as courtship songs at each distance from the female (n=14 birds, r=−0.8552, p=3.07*10^−22^, Pearson’s correlation). Gray lines represent individual birds. Black squares and whiskers represent mean and SEM across all birds for each distance. UN represents songs produced in the undirected condition when the bird was presented with an empty cage.

### Song syllable amplitude became undirected-like with distance

To characterize amplitude independent of the bird’s position within the cage, we used head-attached microphones in a subset of birds (n=7/14 birds, Fig. 2A, Fig. 2B) and examined song syllable amplitude in birds with ≥ 3 courtship song bouts at ≥ 3 distances including L0 (n=5/7 birds). Similar to Brumm and Slater (Brumm and Slater, 2006), some syllables (5/13) increased in amplitude as distance from the female increased (Fig. 2C). However, a similar number of syllables (4/13) decreased in amplitude (Fig. 2D). Further, we found that some syllables (Fig. 2E, red symbols, n=5/13) were louder during undirected song compared to courtship song at the closest distance (referred to as L0 song), while a similar number of syllables were louder during L0 song (Fig. 2E, blue symbols, n=6/13). Interestingly, most syllables that increased in amplitude (positive slopes, red symbols in Fig. 2F) were louder in undirected song compared to L0 song (Fig. 2F, n=4/5) and syllables that decreased in amplitude (negative slopes, blue symbols in Fig. 2F) were louder in L0 song compared to undirected song (Fig. 2F, n=4/4). Irrespective of the actual direction of change, this suggested that syllable amplitudes became more “undirected-like” with increasing distance. This was also true for group data as well after we inverted data from syllables that were louder during undirected song (see Methods). Mean syllable amplitude across all birds (n=5) became more undirected-like with increasing distance (Fig. 2G, Supp. Fig. 2.1 shows the same result averaged across all syllables).

**FIGURE 2.**
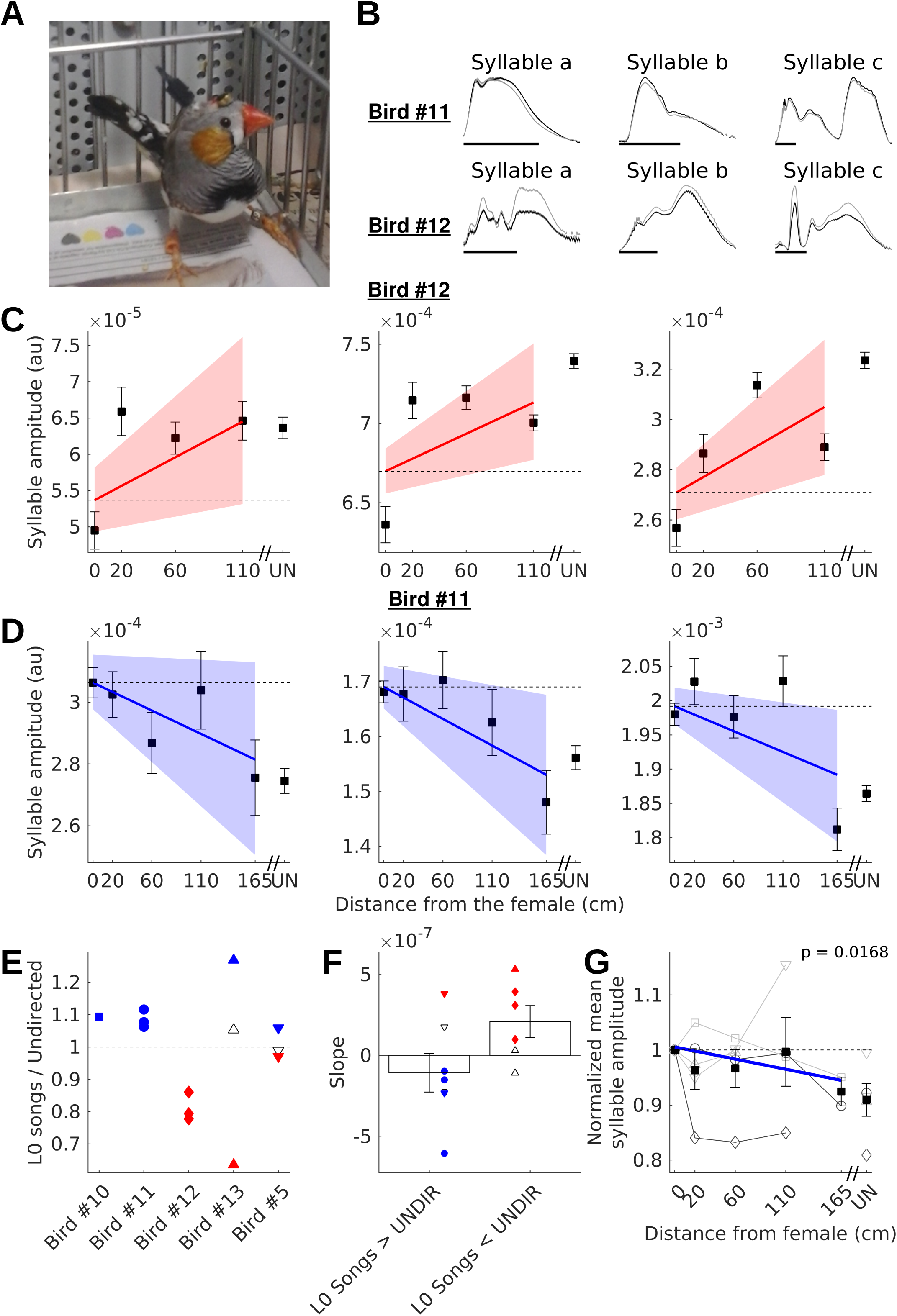
Courtship song ampitude became undirected-like with distance (A) Picture of a male zebra finch (Bird #1) with a head-fixed microphone that was used to record song amplitude independent of the position of the bird within the cage (B) Mean amplitude for syllables for two birds for courtship songs at the shortest distance (L0 songs – shown in black) and for the undirected condition in presence of the empty cage (shown in gray). Shading in all cases is SEM. Scale bar in all plots = 25ms. (C) and (D) show changes in amplitude of song syllables for two example birds (3 syllables each). Each plot shows data for one syllable. In each plot, squares and whiskers represent the mean and SEM for amplitude of the syllable during directed song produced at difference distances from the female. Squares and whiskers corresponding to UN represent mean and SEM for amplitude of the syllable during undirected song produced in the presence of an empty cage. Solid lines represent linear fits to the data. Blue lines represent significant negative slopes, red lines represent significant positive slopes and black lines represent non-significant fits. The shaded region represents 95% confidence intervals estimated using a permutation test (see Methods for details). Dashed line is a line with zero slope representing no relationship between distance and feature value. (E) Ratio of amplitudes for individual syllables for courtship songs at the closest distance (L0) relative to undirected songs is shown. Individual circles represent individual syllables. Symbols for a given bird are used for the same bird in (F) and (G). Red filled symbols represent syllables that are significantly louder during undirected songs, blue filled symbols represent syllables that are significantly louder during L0 songs and black unfilled symbols are syllables that are not significantly different between the two conditions (unpaired t-test for each syllable). (F) shows the slopes for the linear fits for distance dependent changes in syllable amplitude plotted separately for cases where syllable amplitude for L0 DIR (directed song at L0) was greater than UNDIR (undirected song in the presence of an empty cage) and vice versa. Bars and whiskers represent mean and SEM across birds. Blue filled circles represent significant negative slopes and red filled circles represent significant positive slopes. Diamonds represent the bird shown in (D) and circles represent the bird shown in (C). (G) Group data (n=5 birds) with each bird normalized to the feature value for L0 DIR. Symbols and gray lines represent mean syllable amplitude for individual birds. Birds with undirected song amplitude greater than L0-DIR syllable amplitude have been flipped. Darker gray lines represent birds shown in (C) and (D). Squares and whiskers represent mean and SEM across all birds for each distance. Solid lines represent linear fits to the data. Blue lines represent significant negative slopes and black lines represent no significant relationship.

### Other song acoustic features became undirected-like with distance

To examine whether other acoustic properties of courtship songs also became undirected-like with distance, we considered all birds with ≥ 3 directed song bouts at ≥ 3 distances including L0 (n=11/14 birds) and quantified 3 different features of courtship song bouts, namely, (1) number of introductory notes (INs) at the start of each song bout, (2) number of complete song motifs / bout and (3) average motif duration (Fig. 1A). Consistent with earlier studies (Kao et al., 2005; Sossinka and Böhner, 1980), we found significant differences in subsets of birds for all of these features between L0 songs and undirected songs produced in the absence of the female (Supp. Fig. 3.1).

**FIGURE 3.**
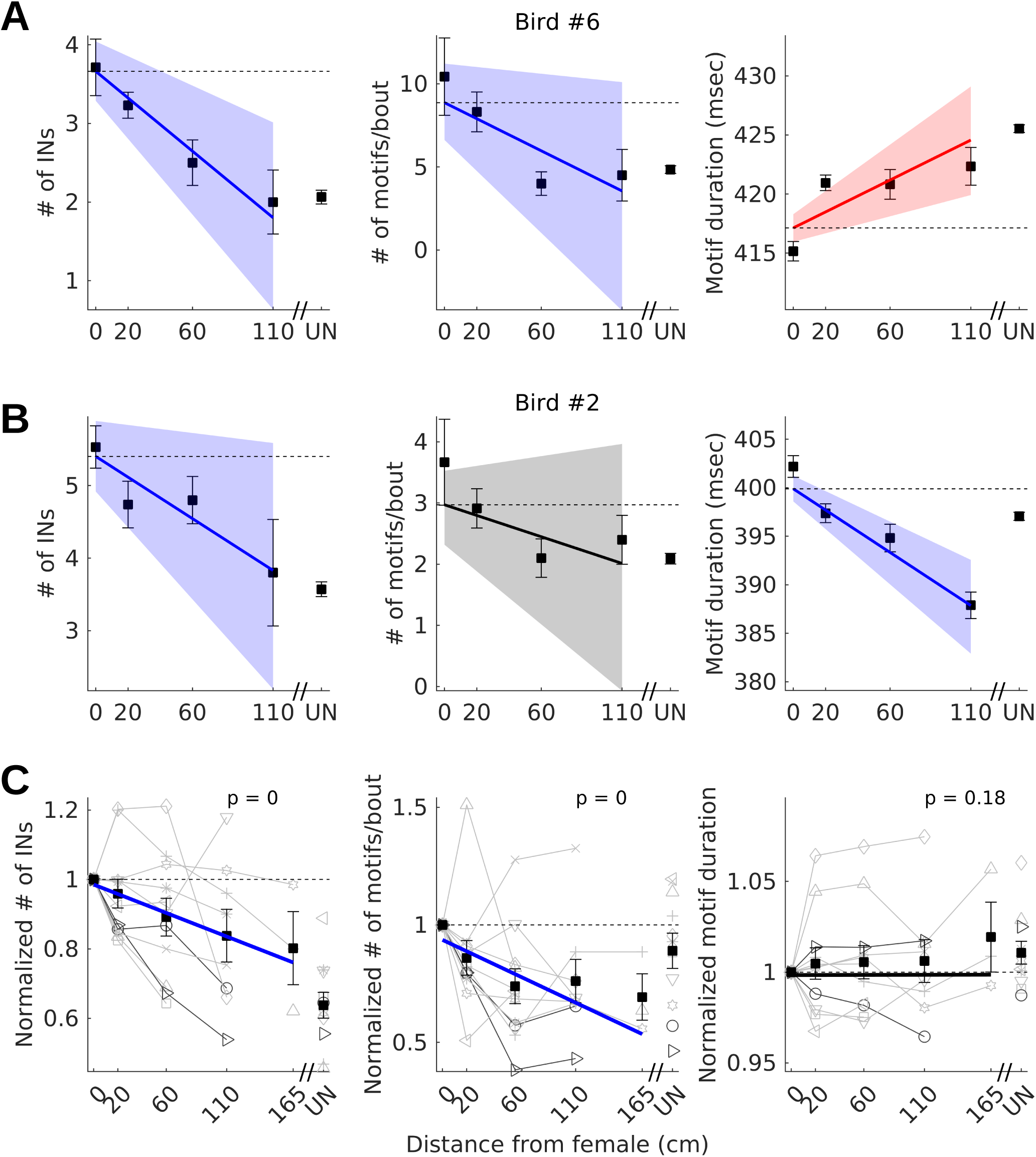
Courtship song features became more undirected-like with increasing distance (A) and (B) show changes in directed song features for two example birds. Left side plot – mean # of INs, middle – mean # of motifs/bout, right – mean motif duration. In each plot, squares and whiskers represent the mean and SEM for directed song features produced at difference distances from the female. Squares and whiskers corresponding to UN represent mean and SEM for undirected song features produced in the presence of an empty cage. Solid lines represent linear fits to the data. Blue lines represent significant negative slopes, red lines represent significant positive slopes and black lines represent non-significant fits. The shaded region represents 95% confidence intervals estimated using a permutation test (see Methods for details). Dashed line is a line with zero slope representing no relationship between distance and feature value. (C) Group data (n=11 birds) with each bird normalized to the feature value for L0 DIR. Symbols and gray lines represent data from individual birds. Left - # of INs, Middle - # of motifs/bout, Right – motif duration. Darker gray lines represent birds shown in (A) and (B). Squares and whiskers represent mean and SEM across all birds for each distance. Solid lines represent linear fits to the data. Blue lines represent significant negative slopes and black lines represent no significant relationship.

All of these features also changed significantly with distance in subsets of birds and became more similar to undirected song features. Similar to the amplitude data, positive slopes (increasing feature value) was mostly associated with higher feature values in undirected songs while negative slopes were mostly associated with higher feature values in L0 songs (Supp. Fig. 3.2) Group data for 2 of 3 features also showed a distance dependent decrease towards undirected feature values (Fig. 3C). Together with the distance-dependent changes in syllable amplitude, these results suggested a change to undirected-like song properties as distance from the female increased.

To control for potential biases introduced by video-based classification of songs, we performed a control analyses by relaxing the criterion for courtship songs (see Methods). This analysis yielded similar results (Supp. Fig. 3.3) further strengthening our conclusions that courtship song features including song amplitude became more undirected-like with increasing distance from the female.

### Female response modulates number of motifs/bout and syllable amplitude

To understand possible causes for this change to undirected-like properties, we examined female response as it is known to modulate courtship song properties (Garson et al., 1980). We scored videos for previously characterized displays corresponding to female responses (Morris, 1954; Zann, 1996) (n=8 birds, primarily L0 and L1, see Methods) and found a significant decrease in female responses as distance increased (Fig. 4A). To see whether the presence or absence of female responses affected song features, we examined cases where males produced ≥ 3 courtship song bouts both with and without female responses at 1 or more distances. Consistent with earlier studies (Garson et al., 1980), the number of motifs/bout was significantly less (undirected-like) when females did not respond (Fig. 4C). Number of INs and motif duration were not significantly affected (Fig. 4B, 4D). In birds with head-attached microphones, song syllables were louder when females responded, though the difference was significant only in 40% of the syllables (n=4/10, p<0.05, unpaired t-test, Fig. 4E). Overall, these data suggested that reduced female response at farther distances is one factor that might contribute to distance-dependent changes in courtship song properties.

**FIGURE 4.**
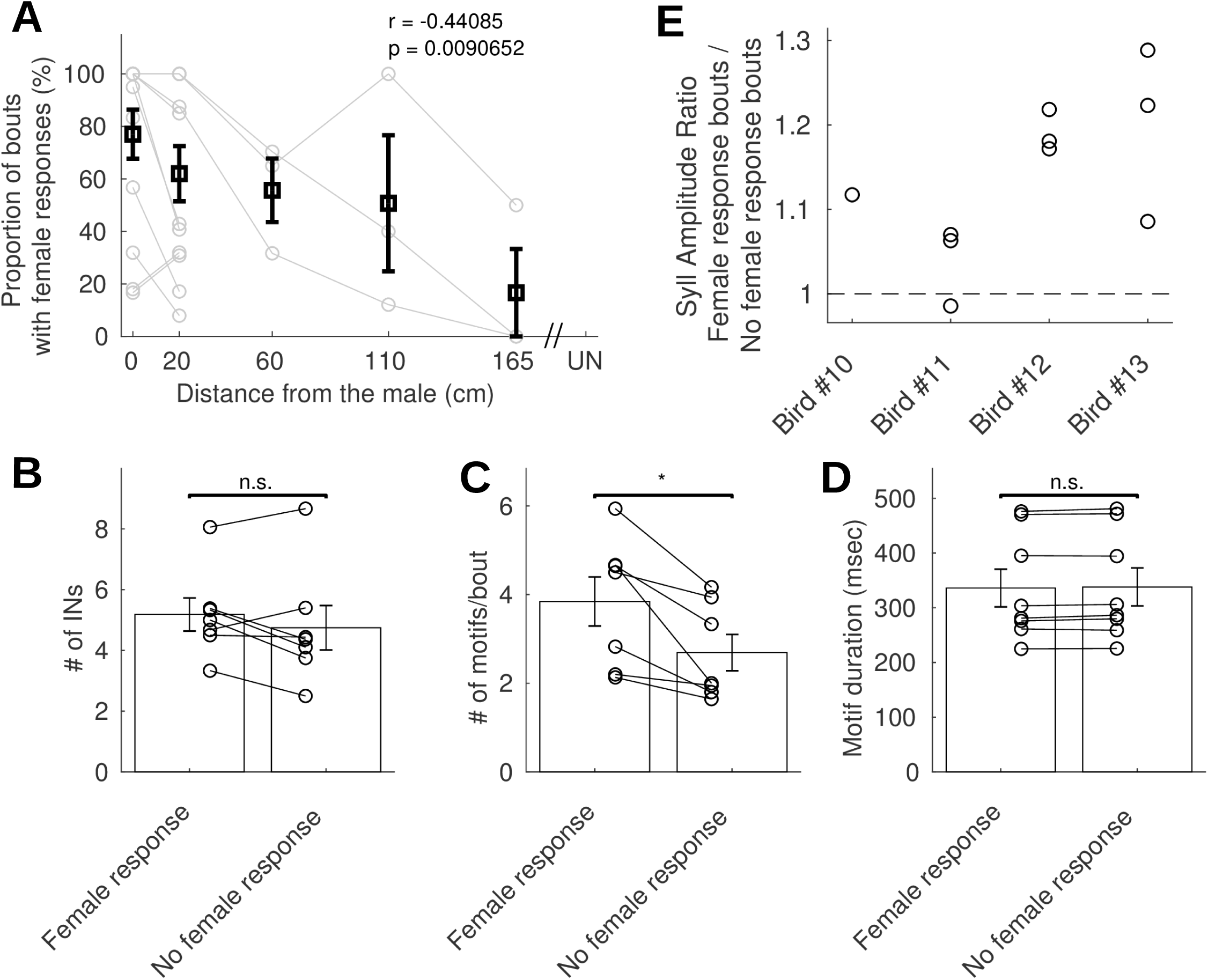
Female response changed with distance and affected courtship song features (A) shows changes in the percentage of courtship song bouts with female responses as distance from the female (n=10 birds, r=−0.44085, p=0.0090652, Pearson’s correlation). Gray lines represent individual birds. Black squares and whiskers represent mean and SEM across all birds for each distance. UN represents songs produced in the undirected condition. (B), (C) and (D) show differences in three song bout features – mean # of INs at the start of a bout (B), # of motifs/bout (C), and mean motif duration (D) - between courtship song bouts with and without female responses. Circles represent individual birds. Bars and whiskers represent mean and SEM across all birds (n=8). * represents p < 0.05. (E) Ratio of syllable amplitude for courtship song bouts with and without female responses. Individual circles represent individual syllables. Dashed line indicates equal syllable amplitude in both types of bouts.

## DISCUSSION

In this study, we examined changes in courtship song amplitude and other acoustic features as distance from a female bird increased. Our results showed that song amplitude and other acoustic features of courtship songs approached corresponding undirected song values with increasing distance from the female. Further, we showed that female responses to song decreased with increasing distance and both syllable amplitude and the number of songs/bout were reduced in the absence of female responses. Overall, these results support the hypothesis that distance-dependent changes in amplitude are part of a more general change in song features towards “undirected-like” song features.

These results suggest that courtship and undirected song properties are two extremes of a continuum of song states, rather than two distinct states (Kao and Brainard, 2006; Kao et al., 2005; Sossinka and Böhner, 1980). Similar graded changes were observed in courtship song properties with changes in the quality of the stimulus triggering song (Bischof et al., 1981). Further, graded changes, with increasing distance from a receiver, have also been shown to occur for aggressive responses of green frogs and claw-waving displays of fiddler crabs (How et al., 2008; Owen and Gordon, 2005). Along with our results, this suggests the possibility that a continuum of behavioral states may be more common than distinct behavioral states.

Overall, our results provide a simple behavioral mechanism that explains changes in song amplitude with increasing distance from the female. Increases in amplitude of vocalizations with increasing distance from a receiver have been observed for Drosophila song (Coen et al., 2016), frog vocalizations (Owen and Gordon, 2005), bird song (Brumm and Slater, 2006) and human speech (Johnson et al., 1981; Michael et al., 1995). In the case of humans, it had been proposed that higher cognitive abilities like perspective taking, were involved in the ability to increase speech amplitude. Based on the observation that zebra finches also increase song amplitude with distance, Brumm and Slater proposed that birds and possibly humans, learn to increase their song amplitude as increased song amplitudes are more effective at eliciting female responses with increasing distance (Brumm and Slater, 2006). Using a more accurate method to measure song amplitude, we found both increases and decreases in song amplitude with increasing distance. Both sets of changes were explained by a more general change to undirected-like song properties. Such a change could be a result of varying levels of motivation caused by increasing distance and / or reduced female responses. Our results suggest a simpler mechanism not requiring learning, whereby increasing distance from the female shifts song state changing a number of acoustic properties including amplitude. This could be a general mechanism for explaining distance-dependent changes in vocalization amplitude.

## METHODS

All experiments were performed at IISER, Pune and were approved by the Institutional Animal Ethics Committee (IAEC) in accordance with the guidelines of the Committee for the Purpose of Control and Supervision of Experiments on Animals (CPCSEA), New Delhi.

Fourteen adult male zebra finches and six adult female zebra finches (> 120 days post hatch), used in this study, were either purchased from outside sources (n=8 males and n=2 females) or bred in the bird colony (n=6 males and n=4 females) at IISER, Pune. They were housed in group cages in a colony with a 14 hour (light): 10 hour (dark) cycle. All birds were first separated into individual cages and isolated in sound-attenuating chambers, 7 days prior to the start of the experiment.

### Experimental design

Experiments began on the 7^th^ day after isolation and on each experimental day, the male bird was placed in a cage in the recording room about an hour prior to the start of the experiment. A camera (Logitech Webcam HDC525) was positioned to record videos of movements of the male bird using ffmpeg (https://www.ffmpeg.org/). Songs were recorded using different methods as outlined in the next section. For a subset of males, we also recorded female movements either manually during the experiment (n=8 birds) or using another camera (n=2 birds).

During each recording session, a cage with a female was presented at a specified distance from the end of the male’s cage. Recording sessions continued until birds had produced atleast 10 song bouts (median session duration = 59 minutes; range = 7-273 minutes). On a given day, 1-6 sessions were conducted with 20 minute intervals between consecutive sessions. The distance of the female cage in each session was randomly chosen from one of 5 pre-specified distances (see Fig. 1C: L0 - 0 cm, L1 - 20cm, L2 - 40cm, L3 - 110 cm and L4 - 165 cm). Undirected songs in the absence of a female were recorded in separate sessions by placing an empty cage at a distance of 20cm (at L1). As far as possible, such undirected song sessions were interleaved with other female presentation sessions. However, if birds did not sing enough undirected songs during these sessions, we recorded undirected songs during separate longer sessions soon after the completion of female presentation sessions (median undirected session duration = 127 minutes; range = 26-537 minutes). As the L0 distance was not in our initial experimental design, for a subset of birds (n =5), songs at L0 (0 cm) were recorded separately 2-3 months after the other sessions. Undirected songs were also recorded at the same time. Data from all sessions at a particular distance were combined together as there were no consistent statistical differences between sessions for each bird (Kruskal-Wallis test).

### Song Recordings

Songs were recorded using three different methods.

1. <u>Mixer microphone (MM)</u> - A lavalier microphone (AKG C417PP) was connected to a mixer (Behringer XENYX 802) and placed at a fixed position on the male bird’s cage throughout the recording session. This method showed considerable variability in song amplitude within and across sessions (Supp. Fig. 5A). Such recordings were carried out in n=5 birds and were included for analysis of song features excluding amplitude and FF.
2. <u>Backpack Microphone (BPMic)</u> – We adapted a previously described microphone backpack design (Anisimov et al., 2014) with a small microphone (CUI, CME-1538-100LB purchased from Mouser Electronics, TX). This was attached to the back of the bird with the help of a thread (Total weight = 0.89g). Birds were gently restrained and the backpack was attached in a short procedure that lasted a few minutes. Birds took a few days (3-7 days) to resume singing after backpacks had been fixed. Syllable amplitudes recorded by this method still exhibited considerable variation, albeit less than the mixer microphone. Such recordings were done in n=8 birds and this data was used for measurement of other song features except amplitude.
3. <u>Head Fixed Microphone (HFMic)</u> - To record song amplitude accurately, we surgically implanted microphones (same as those used in the backpack; CUI, CME-1538-100LB purchased from Mouser Electronics, TX) along with a connector (weighing less than 0.5g) on the bird’s head. Surgical procedures used were similar to previous studies that have implanted microdrives with electrodes for electrophysiological recordings (Okubo et al., 2014). Briefly, birds were given an analgesic (meloxicam) two hours prior to the start of surgery and were subsequently anesthetised using ketamine (30mg/kg), xylazine (1mg/kg), and diazepam (7mg/kg). A local anesthetic (lidocaine – 2%; 50-75μl) was injected sub-cutaneously before making an incision. A small piece (5 mm x 5 mm) of skin was removed from the top of the head and holes were made in the top layer of skull. The microphone with connector was placed on the inner layer of skull and cemented to the skull using dental cement (Panavia F2.0). Song recordings with head-attached microphones had the lowest within and across session variability and were used for analysis of changes in song amplitude with distance (Supp Fig.5A).

In all cases, the output (mixer or backpack / head-fixed microphone) was directly connected to a sound card on the computer and data was recorded using custom-written software in Python. We recorded data in 30 second files continuously for each session. However, we noticed that occasionally, some data (< 0.5s) was lost between two consecutive files. Therefore, we excluded all song bouts that extended between two consecutive files.

For one of the birds, we recorded data at different time-points with all 3 types of microphones and all of the data was combined for analysis of all song features except amplitude. For amplitude, we only used recordings with the head-fixed microphone. For 4 birds, we recorded data at different time-points with 2 types of microphones (MM and BPMic) and all data was combined for analysis of song features.

### Video Analysis of male displays during song

During each recording session, song bouts were first identified in the audio recordings. The corresponding portions of the videos were later extracted for video scoring. We noted whether the male was looking in the direction of the female or not and we also scored the videos for the presence or absence of 4 different movements (puffing up, beak-wipes, hops, and turn-arounds) that have been previously documented to be present during directed singing (Morris, 1954; Ullrich et al., 2016; Williams, 2001; Zann, 1996). Song bouts were then classified into one of 3 categories as follows:

- Directed (D) - if the male looked in the direction of the female and performed any two of the above mentioned movements
- Ambiguous (AMBIG) - if the male only looked in the direction of the female but did not perform any of the movements
- Undirected (UN) - if the male looked away from the female

Same scoring scheme was also used to score videos of song bouts produced in the absence of female.

To validate our video scoring scheme, we had three other observers score a random subset of videos (n=90 song videos). These observers were familiar with the zebra finch model system and were provided information about the scoring criteria outlined above, but were blind to the context in which songs were produced. Overall, for all songs scored, at least two of the observers agreed with the experimenter 77% (69/90) of the cases (Supp. Fig. 1A). This provided considerable validation for the scoring scheme employed by the experimenter.

In addition to this validation, we also performed a control analyses to check for the influence of video scoring on the results. Both D and AMBIG songs were considered as courtship songs and this constituted a relaxation of the criterion for courtship songs.

### Video analysis of female responses to song

In each video associated with a song bout, vocalizations and movements of the female bird (puffing up, hops, tail quivering) along with her position with respect to the male bird were noted. Based on these, female responses were divided into 3 categories

- Responding (R) - If the female was close to the male, looked in his direction and performed any one of the above mentioned movements or called in response to song.
- Ambiguous (AMBIG) – if the female was close to the male and looked in his direction, but did not call or perform any movements
- Not responding (NR) – if the female was far away from the male and did not look in his direction

Only the R category was considered as a response and used for further analysis.

### Song analysis

Custom-written MATLAB code was used for all analysis. Files with vocalizations for each session were identified through an initial screening process and vocalisations were segmented based on an amplitude threshold. This amplitude threshold was chosen separately for each bird based on visual inspection of a few song files and maintained constant across all days and sessions for a given bird. Syllables with inter-syllable intervals < 5ms were merged and syllables that were shorter than 10ms were discarded. Individual syllable types were identified visually and one template was made for each of the different syllable types. Syllables in all files were first labelled using a template matching procedure (adapted from (Glaze and Troyer, 2007)) and then checked manually for consistency.

We identified a few types of non-song vocalizations (short calls, long calls, intermediate calls) and two types of song elements – introductory notes, and motif syllables. Short repeated vocalisations occurring at the beginning of song bouts were labelled introductory notes (INs). Stereotyped sequences of syllables repeated in a continuous manner or interrupted by calls or INs were termed as motifs. Groups of vocalizations with at least one motif element were segregated into song bouts based on the presence of 2s of silence before and after the group of vocalizations.

To characterize differences in song features with distance, we measured 5 different features.

- Number of INs at the beginning of a bout – Starting backwards from the beginning of the first motif syllable in a bout, we counted all consecutive introductory notes that were separated by < 500ms (Kao and Brainard, 2006; Rajan and Doupe, 2013; Stepanek and Doupe, 2010).
- Number of motifs / bout – For each bird, we first identified the most commonly occurring motif type (only 1 motif type for n=13 birds and 2 motif types for n=1 bird) and counted the number of complete motifs of this type for each bout (Sossinka and Böhner, 1980).
- Motif Duration – As a measure of tempo of song, we considered the duration of the most commonly occurring motif type for each bird as the time between offset of the last syllable of the motif and onset of the first syllable of the motif. We considered only complete motifs and calculated average motif duration across all complete motifs at a particular distance.
- Log Amplitude - Log amplitude was as described previously (Kao et al., 2005). Briefly, the raw sound waveform was bandpass filtered between 300 and 8000 Hz, rectified and smoothed with a square filter of length 2ms. Log amplitude was then calculated as the the area under this curve between syllable onset and offset. Log amplitude was measured for all motif syllables and averaged across all occurrences of the syllable at each distance within courtship song bouts.

### Assessment of changes in song features with distance

Birds with ≥ 3 courtship song bouts at ≥ 3 distances including at the closest distance L0 were considered for analysis. For each feature, parameters of a linear fit to data from individual bouts across distances was estimated using the Matlab function robustfit to minimize the influence of outliers. The significance of the fit was assessed by using a bootstrap procedure to calculate the 95% confidence intervals for the slope and intercept. Briefly, the residuals of the fit were randomly resampled and 2000 new sets of data were generated. robustfit was again used to calculate parameters of a linear fit for these 2000 data sets and 95% confidence intervals for the slope and intercept parameters were calculated. We considered the slope as significant if the 95% confidence interval did not include 0 (zero slope would indicate no relationship between distance and feature value). At the individual bird level, the slope was used as a measure of the direction of change.

At the group level, we normalized feature values for each bird by the feature value at the closest distance (L0). Across all birds, we did not have data for all of the distances since different birds sang at difference subsets of distances. To assess significance of the change at the group level, we used a procedure that has been used for datasets like ours which are characterized by greater inter-individual variation and missing values at random locations (Brumm and Slater, 2006; Mundry, 1999). Briefly, the procedure involved resampling the data within each bird while maintaining the combination of distances with data. For eg. if we had data for one bird at distances L0, L2 and L3, then the feature values at these distances were resampled to generate a new set of values for L0, L2 and L3 while maintaining no values at L1 and L4. This way, we generated 10000 surrogate datasets for the group data and used robustfit to calculate parameters of a linear fit to the group data. The actual slope value was considered significant if the probability of observing a slope as high or as low as the actual slope value was < 0.05.

For amplitude, we carried out an extra step to remove outliers before fitting the data. In both cases, for each syllable, outliers were first removed based on duration and then based on amplitude. Values < 3 * inter-quartile range below the 25^th^ percentile of data across all distances or > 3 * inter-quartile range above the 75^th^ percentile of data were considered outliers. Visual inspection of these outliers showed that they were mostly due to mislabelled syllables or syllables with other sources of noise. In most cases, < 1% of syllables were removed as outliers (range 0 – 5.5%).

### Statistical Analysis

All the tests used in this study were carried out with a 95% confidence interval (alpha = 0.05).

To test the significance of variation in number of directed songs with distance, correlation with distance was calculated using Pearson’s correlation test (matlab function corr).

For the comparison of feature differences between L0 songs and ALONE songs, at the individual bird level, we used Kruskal-wallis test (matlab function kruskalwallis) for comparing number of INs and number of motifs, unpaired t-test (matlab function ttest2) for comparing motif duration and log amplitude and an F-test (matlab function vartest2) for comparing variance of FF for individual motif syllables. Significance of group data across all birds for context-dependent changes was tested using the Wilcoxon paired sign-rank test (matlab function signrank).

## ACKNOWLEDGMENTS

We thank Divya Rao for help with surgery and Annika Heinze, Aarcha Thadi, Aboli Ektare for help with recording and labelling songs. We thank Divya Rao, Shikha Kalra, and Aditi Agarwal for scoring videos, Deepak Barua for help with statistical analysis and Aurnab Ghose for helpful discussions. We also thank Girish Deshpande, Deepa Subramanyam, Mimi Kao, Hamish Mehaffey, Anand Krishnan, Divya Rao, and Shikha Kalra for their comments on the manuscript.

## AUTHOR CONTRIBUTIONS

HS and RR designed the experiments, analyzed the data and wrote the paper. HS did all the experiments.

## SUPPLEMENTARY FIGURE LEGENDS

**SUPPLEMENTARY FIGURE 1.1.**
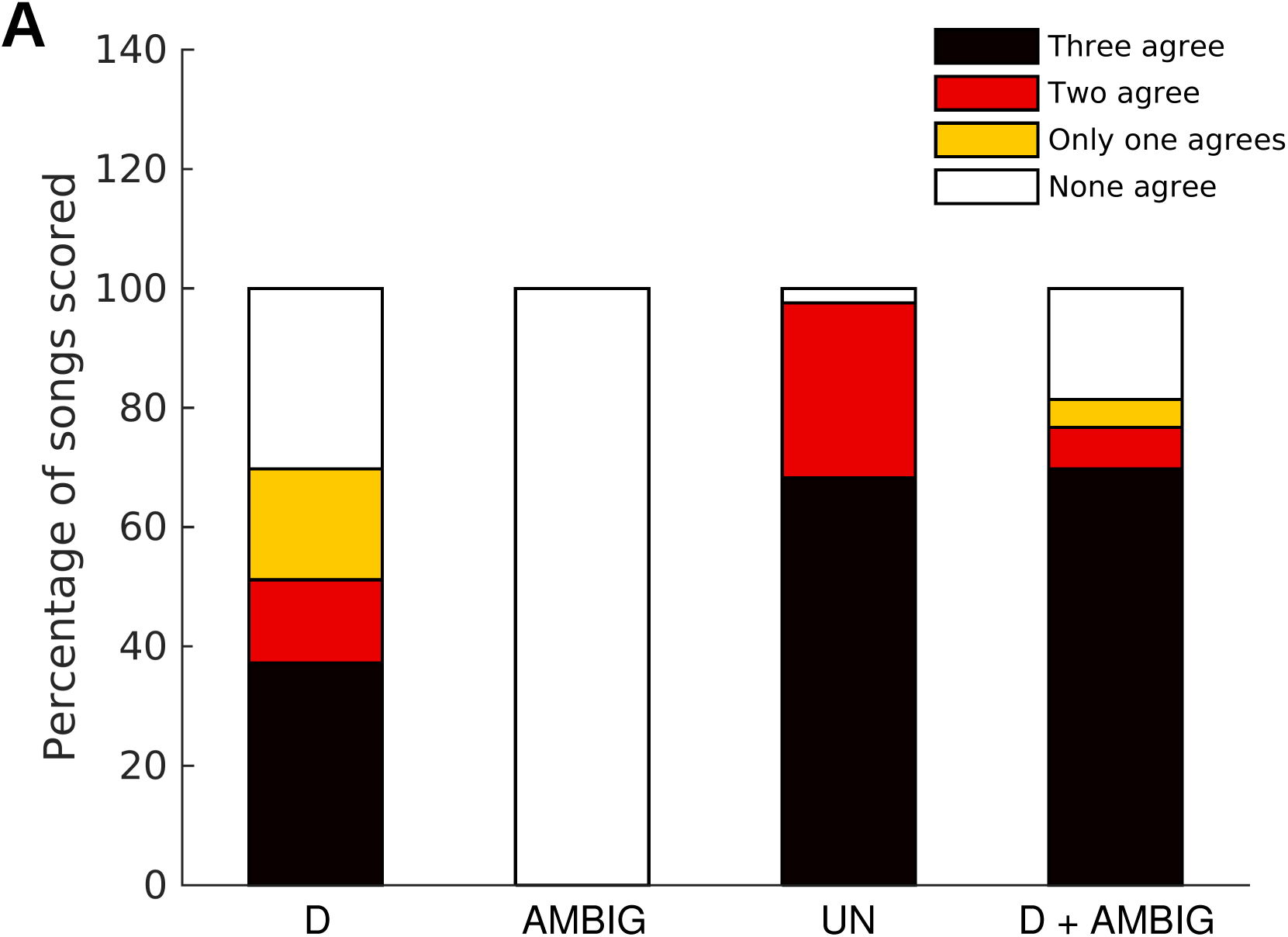
Validation of video scoring (A) Bar graph depicting agreement between video scoring of song bouts between three observers blind to the experimental condition and the experimenter. The x-axis represents categories scored by the experimenter. For the last category, all of songs scored as D by the experimenter were checked for agreement while considering both D and AMBIG scoring by 3 observers.

**SUPPLEMENTARY FIGURE 2.1.**
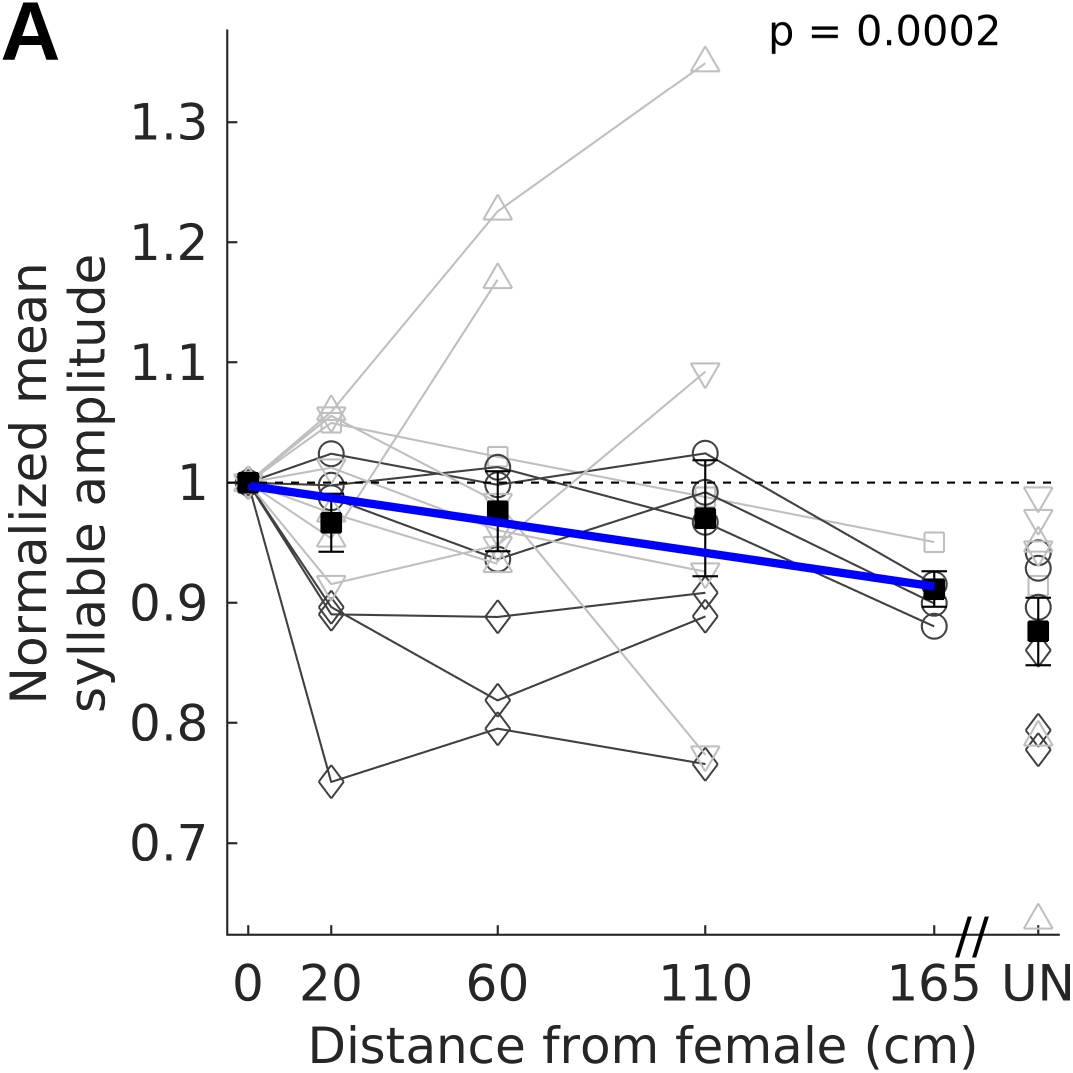
Courtship song syllable ampitude became undirected-like with distance (A) Group data (n=13 syllables from 5 birds) with syllable normalized to the amplitude for L0 DIR. Symbols and gray lines represent mean syllable amplitude for individual syllables. Syllables with undirected song amplitude greater than L0-DIR syllable amplitude have been flipped. Darker gray lines represent syllables shown in 5A and 5B. Squares and whiskers represent mean and SEM across all syllables for each distance. Solid blue line represents linear fit to the data with significant negative slopes (p < 0.05, Permutation test, see Methods).

**SUPPLEMENTARY FIGURE 3.1.**
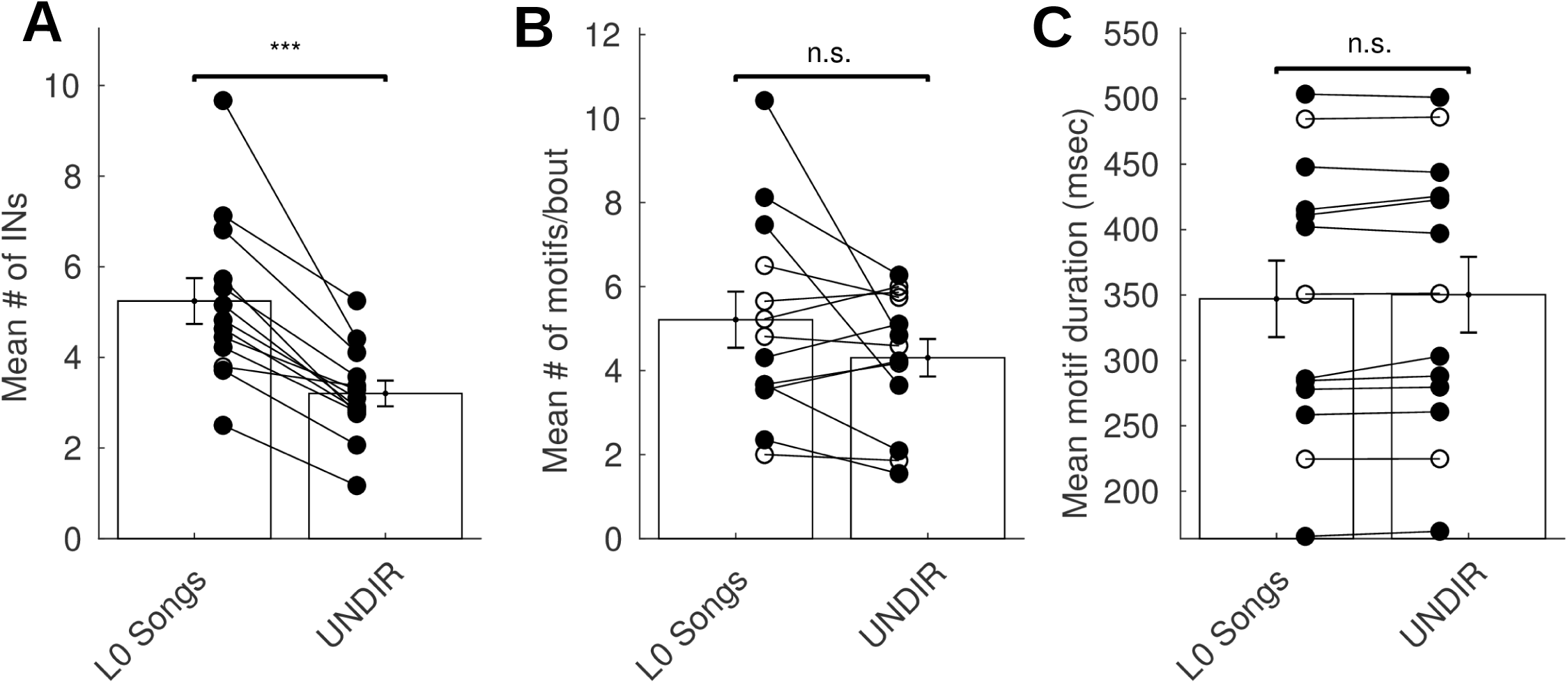
L0 song bouts were preceded by more INs, had more motifs and were longer than UNDIR song bouts (A), (B), and (C) show differences in three song bout features – mean # of INs at the start of a bout (A), # of motifs/bout (B), and mean motif duration (C) - between L0 songs (courtship songs at L0) and UNDIR song bouts (undirected song bouts produced in the absence of the female). Filled circles represent individual birds that have significant differences and open circles represent individual birds with non-significant differences (Kruskal-wallis test for # of INs and # of motifs/bout; unpaired t-test for motif duration). Bars and whiskers represent mean and SEM across all birds (n=13). *** represents p < 0.001.

**SUPPLEMENTARY FIGURE 3.2.**
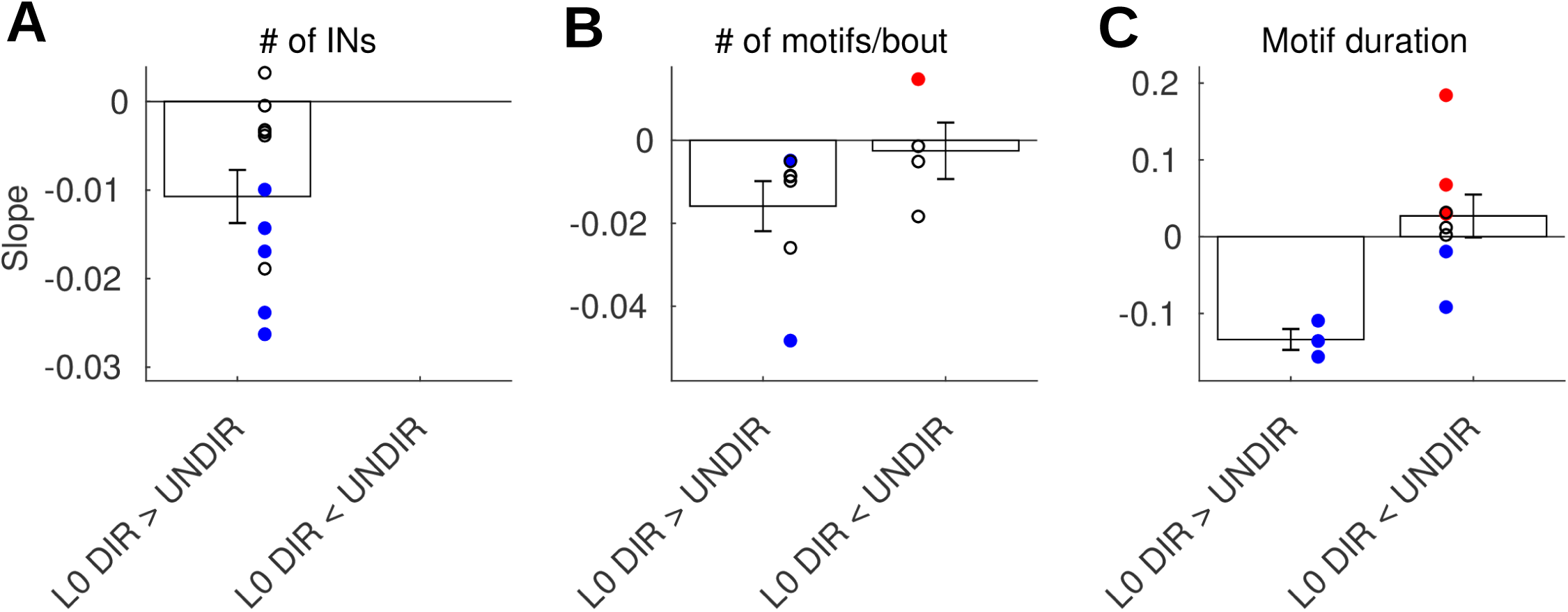
Directed song features became undirected-like with distance (A), (B), and (C) show the slopes for the linear fits for distance dependent changes in different features plotted separately for cases where the feature value for L0 DIR (directed song at L0) is greater than UNDIR (undirected song in the presence of an empty cage) and vice versa. Circles represent individual birds. Bars and whiskers represent mean and SEM across birds. Blue filled circles represent significant negative slopes and red filled circles represent significant positive slopes.

**SUPPLEMENTARY FIGURE 3.3.**
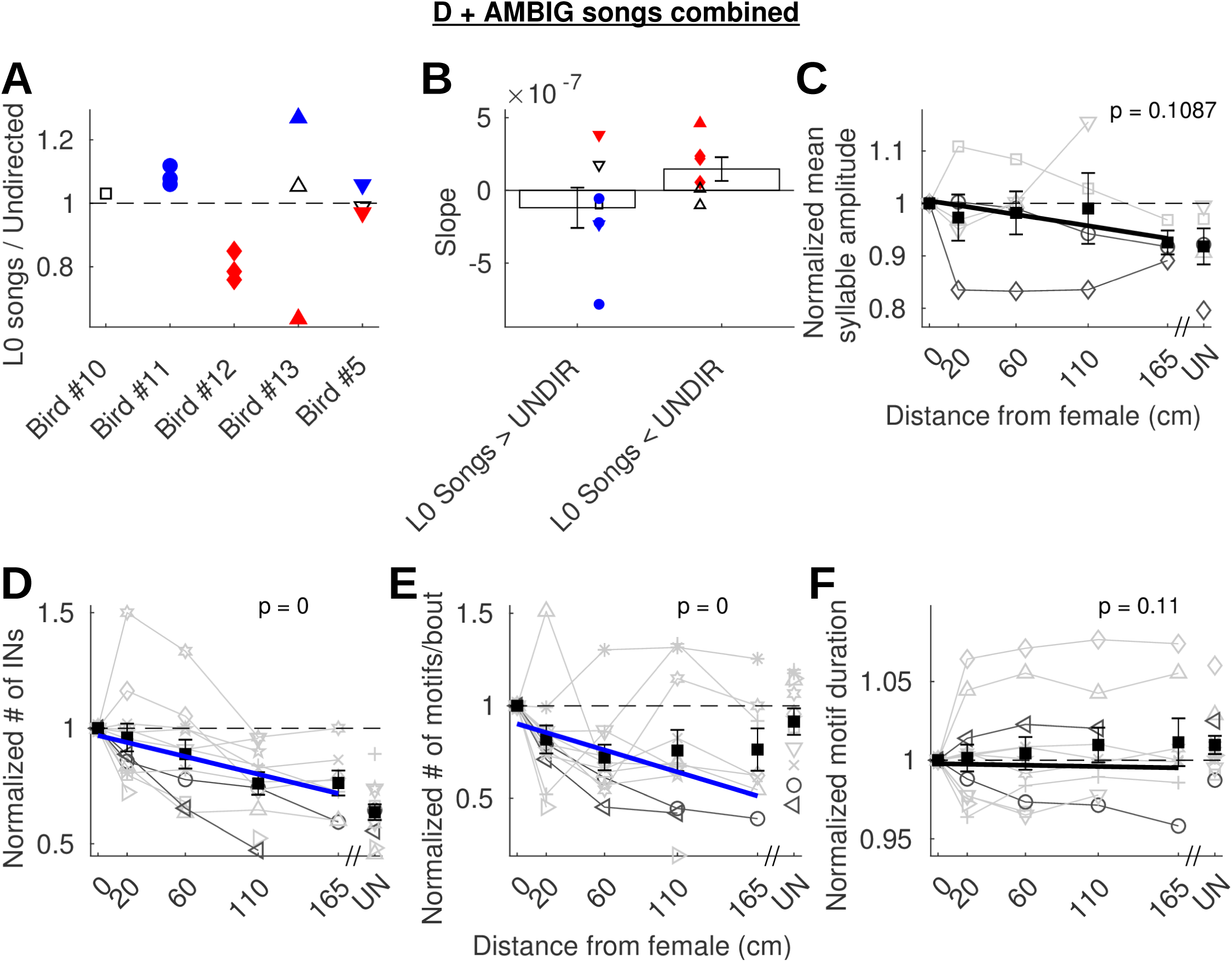
Courtship song features became undirected-like with distance even after criterion for classification of courtship songs was changed to include songs that were ambiguous A. Ratio of amplitudes for individual syllables for courtship songs at the closest distance (L0) relative to undirected songs is shown. Individual circles represent individual syllables. Symbols for a given bird are used for the same bird in (B) and (C). Red filled symbols represent syllables that are significantly louder during undirected songs, blue filled symbols represent syllables that are significantly louder during L0 songs and black unfilled symbols are syllables that are not significantly different between the two conditions (unpaired t-test for each syllable). B. shows the slopes for the linear fits for distance dependent changes in syllable amplitude plotted separately for cases where syllable amplitude for L0 DIR (directed song at L0) was greater than UNDIR (undirected song in the presence of an empty cage) and vice versa. Bars and whiskers represent mean and SEM across birds. Blue filled circles represent significant negative slopes and red filled circles represent significant positive slopes. Diamonds represent the bird shown in (2D) and circles represent the bird shown in (2C). C. Group data (n=5 birds) with each bird normalized to the feature value for L0 DIR. Symbols and gray lines represent mean syllable amplitude for individual birds. Birds with undirected song amplitude greater than L0-DIR syllable amplitude have been flipped. Darker gray lines represent birds shown in (2C) and (2D). Squares and whiskers represent mean and SEM across all birds for each distance. Solid lines represent linear fits to the data. Blue lines represent significant negative slopes and black lines represent no significant relationship. D. (D), (E) and (F) Group data (n=11 birds) with each bird normalized to the feature value for L0 DIR. Symbols and gray lines represent data from individual birds. # of INs (D), # of motifs/bout (E), motif duration (F). Darker gray lines represent birds shown in (3A) and (3B). Squares and whiskers represent mean and SEM across all birds for each distance. Solid lines represent linear fits to the data. Blue lines represent significant negative slopes and black lines represent no significant relationship.

